# Cortico-striatal activity characterizes human safety learning via Pavlovian conditioned inhibition

**DOI:** 10.1101/2021.11.09.467993

**Authors:** Patrick A.F. Laing, Trevor Steward, Christopher G. Davey, Kim L. Felmingham, Miguel Angel Fullana, Bram Vervliet, Matthew D. Greaves, Bradford Moffat, Rebecca K. Glarin, Ben J. Harrison

**Affiliations:** Melbourne Neuropsychiatry Centre, Department of Psychiatry, The University of Melbourne; Melbourne School of Psychological Sciences, The University of Melbourne; Adult Psychiatry and Psychology Department, Institute of Neurosciences, Hospital Clinic, Barcelona, Spain; Institut d’Investigacions Biomèdiques August Pi i Sunyer (IDIBAPS), Centro de Investigación Biomédica en Red de Salud Mental (CIBERSAM), Barcelona, Spain; Laboratory of Biological Psychology, Faculty of Psychology and Educational Sciences, KU Leuven, Belgium; Leuven Brain Institute, KU Leuven, Belgium; The Melbourne Brain Centre Imaging Unit, Department of Radiology, The University of Melbourne

**Keywords:** conditioned inhibition, safety learning, prediction error, dorsal striatum, vmPFC, 7-Tesla fMRI

## Abstract

Safety learning generates associative links between neutral stimuli and the absence of threat, promoting the inhibition of fear and security-seeking behaviours. Precisely how safety learning is mediated at the level of underlying brain systems, particularly in humans, remains unclear. Here, we integrated a novel Pavlovian conditioned inhibition task with ultra-high field (UHF) fMRI to examine the neural basis of inhibitory safety learning in 49 healthy participants. In our task, participants were conditioned to two safety signals: a conditioned inhibitor that predicted threat-omission when paired with a known threat signal (A+/AX-), and a standard safety signal that generally predicted threat-omission (BC-). Both safety signals evoked equivalent autonomic and subjective learning responses but diverged strongly in terms of underlying brain activation. The conditioned inhibitor was characterized by more prominent activation of the dorsal striatum, anterior insular and dorsolateral prefrontal cortex compared to the standard safety signal, whereas the latter evoked greater activation of the ventromedial prefrontal cortex, posterior cingulate and hippocampus, among other regions. Further analyses of the conditioned inhibitor indicated that its initial learning was characterized by consistent engagement of dorsal striatal, midbrain, thalamic, premotor, and prefrontal subregions. These findings suggest that safety learning via conditioned inhibition involves a distributed cortico-striatal circuitry, separable from broader cortical regions involved with processing standard safety signals (e.g., CS-). This cortico-striatal system could represent a novel neural substrate of safety *learning*, underlying the initial generation of ‘stimulus-safety’ associations, distinct from wider cortical correlates of safety processing, which facilitate the behavioral *outcomes* of learning.

**Significance statement:** Identifying safety is critical for maintaining adaptive levels of anxiety, but the neural mechanisms of human safety learning remain unclear. Using ultra-high field fMRI, we compared learning-related brain activity for a conditioned inhibitor, which actively predicted threat-omission, and a standard safety signal (CS-), which was passively unpaired with threat. The inhibitor engaged an extended circuitry primarily featuring the dorsal striatum, along with thalamic, midbrain, and premotor/prefrontal cortex regions. The CS-exclusively involved cortical safety-related regions observed in basic safety conditioning, such as the vmPFC. These findings extend current models to include learning-specific mechanisms for encoding stimulus-safety associations, which might be distinguished from expression-related cortical mechanisms. These insights may suggest novel avenues for targeting dysfunctional safety learning in psychopathology.

## Introduction

Safety learning builds associations between neutral stimuli and the absence of threat, facilitating the inhibition of fear in safe situations. Impaired safety learning is thought to contribute to the pathophysiology of anxiety-related disorders (Grasser and Jovanovic, 2021; van Rooij and Jovanovic, 2019), leading to a renewed interest in its brain-behavioral basis across species (Fendt et al., 2021). Despite its compelling clinical relevance, safety learning remains understudied in humans, and presents several conceptual and empirical challenges (Laing and Harrison, 2021). For instance, while safety signals elicit fewer threat responses compared to fear signals, the same occurs for neutral or ambiguous stimuli, which predict neither the presence or absence of threat (Rescorla, 1969). Neurobiological studies suggest, however, that safety is not a neutral state, but one that conveys information critical to survival and well-being (Grupe and Nitschke, 2013; Tashjian et al., 2021).

Prevailing neuroimaging evidence for human safety learning stems from differential fear conditioning studies, where a standard safety signal (CS-) is compared to a conditioned threat stimulus (CS+). These studies consistently identify increased activity of the ventromedial prefrontal cortex (vmPFC), posterior cingulate cortex, and hippocampus in discriminating CS-from CS+ (Fullana et al., 2015). Though robust, it is unclear to what extent these findings directly reflect safety learning *per se*. For example, animal models demonstrate a role for the vmPFC in expressing fear inhibition at test, but less involvement during initial learning (Gewirtz et al., 1997; Kreutzmann et al., 2020; Sarlitto et al., 2018). Further, while medial prefrontal cortical pathways from the ventral tegmentum and hippocampus facilitate the post-conditioning use of safety information (Meyer et al., 2019; Yan et al., 2019), regions such as the insular cortex and dorsal striatum (caudate and putamen) show learning-specific involvement during conditioning (Christianson et al., 2008; Christianson et al., 2011; Foilb et al., 2016; Rogan et al., 2005). These findings suggest that important differences exist between brain systems that acquire safety information via conditioning and those that subsequently express this information in the form of fear inhibition and affective appraisal (Battaglia et al., 2021; Tashjian et al., 2021).

In order to isolate learning-specific mechanisms in the brain, human studies could integrate paradigms more directly informed by fundamental principles from associative learning theory. We have proposed that the ‘Pavlovian conditioned inhibition’ paradigm leverages these principles to provide an optimal experimental model of safety learning (Laing and Harrison, 2021). In this paradigm, a CS is reinforced alone (A+), but not reinforced when combined with a second CS (AX-). The ‘conditioned inhibitor’ thereby indicates *threat-omission* in proximity to threat signals. Nonreinforcement of ‘AX-’ evokes a salient mismatch between expected threat-delivery and actual threat-omission – inducing a prediction error (Yau and McNally, 2018). Established learning theories predict that this mechanism generates robust links between the inhibitor and threat-omission (Schultz and Dickinson, 2000; Wagner and Rescorla, 1972), providing an operational definition of safety learning (Laing and Harrison, 2021). The paradigm has been shown to produce conditioned safety in human behavioral studies (Laing et al., 2021; Neumann et al., 1997), but is yet to be investigated in neuroimaging.

The aim of the current study was to examine the neural basis of human safety learning via Pavlovian conditioned inhibition combined with ultra-high field fMRI. Our first aim was to directly compare the neural correlates of a conditioned inhibitor, which actively signaled threat omission (A+/AX-), and a standard safety signal, which was passively unreinforced (BC-). Second, we aimed to identify regions underlying the initial learning of ‘stimulus-safety’ associations by contrasting early and late conditioning trials, and comparing responses during conditioning to those during a subsequent test phase. We hypothesized that the conditioned inhibitor would more distinctly engage subcortical circuitry, particularly striatal and midbrain regions, which have well-established roles in prediction error-based reinforcement learning (Garrison et al., 2013; Papalini et al., 2020), whereas the standard safety signal would involve distributed cortical regions, including the vmPFC, linked to the cognitive evaluation of learned safety.

## Material and Methods

### Participants

Forty-nine participants were recruited to the study. All participants met the following eligibility criteria: (i) they were aged between 18-35 years, (ii) had no current or past diagnosis of mental illness as per screening via the Mini-International Neuropsychiatric Interview (MINI, Sheehan et al., 1998), (iii) were fluent in English, (iv) were not taking any psychoactive medications, and (v) had no contraindications to MRI, including pregnancy. All participants had normal or corrected-to-normal vision and provided written informed consent, following a complete description of study protocol, which was approved by The University of Melbourne Research Ethics Committee. Of the initial sample, two participants did not complete scanning (one due to technical failure; one who discontinued), and a further four were excluded due to excessive head motion (see image pre-processing). The final sample consisted of 43 participants (20 female) with a mean age of 24.35 years (± 4.35).

### Safety learning task

Participants completed a novel Pavlovian conditioned inhibition task (Figure 1), adapted from a prior behavioral paradigm (Laing et al., 2021). A series of geometric figures were used as conditioned stimuli (CS) and the aversive unconditioned stimulus (US) was a 95-dB white noise of 500ms duration. Inter-trial-intervals (ITI) were jittered between 8-12s (*M* = 10s) and featured a white fixation cross in the center of a black screen. Each CS had a duration of 5s total, firstly presented for 1s alone, then joined by a threat-expectancy rating scale for a further 4s, after which the US was delivered or omitted. Threat-expectancy ratings were made on a 9-point scale, using a button box in participants’ right hand. The scale displayed the following labels: ‘DEFINITELY NO’ at the left-most end, ‘NOT SURE’ in the center, and ‘DEFINITELY YES’ at the right-most end, and was automatically centered at ‘NOT SURE’ upon each new presentation. To enhance inhibitory learning and support the analysis of safety signal responses, the task included a 100% reinforcement rate for all CS+. This choice ensured that A+ maintained a robust threat association, such that US-omission following AX-would reliably violate participants’ threat expectation (Harris et al., 2014; Laing and Harrison, 2021; Lysle and Fowler, 1985). While partial CS+ reinforcement is typically used to avoid the US confounding of CS+ responses (Fullana et al., 2015), this was unnecessary in the current study, which was designed to target safety signals exclusively.

**Figure 1.**
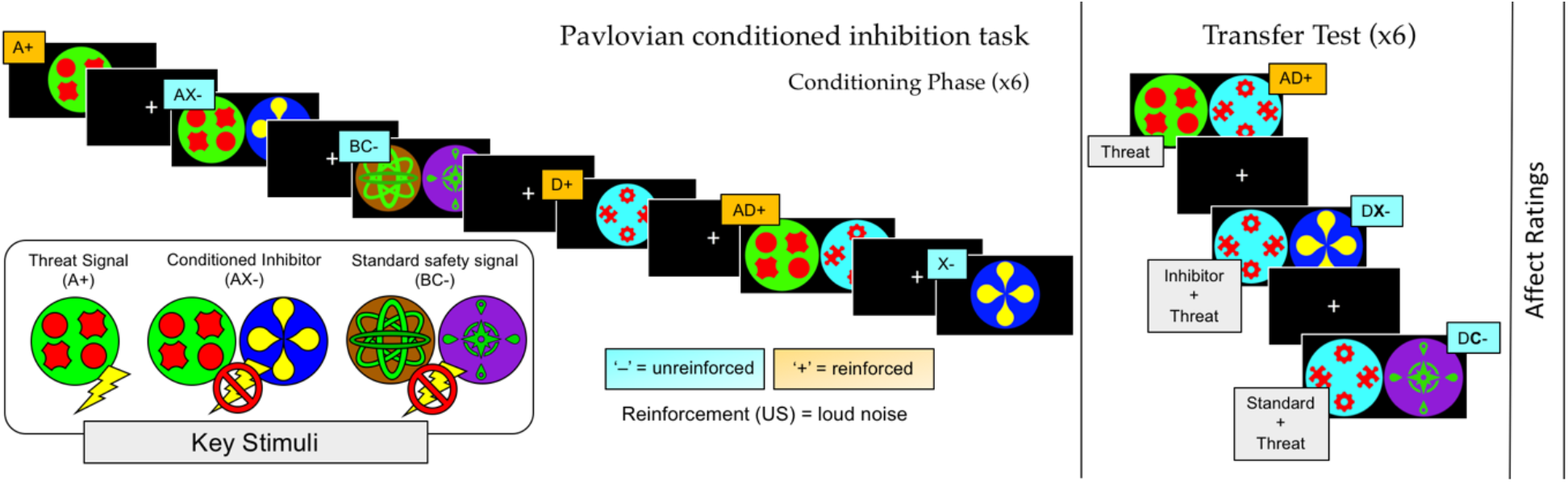
Pavlovian conditioned inhibition fMRI task, adapted from Laing et al. (2021). Six stimulus configurations were presented during conditioning. The transfer test occurred immediately following conditioning and featured the two safety signals (inhibitor ‘X’ and standard ‘C’) combined with a conditioned threat cue (D+). Between each CS trial, the inter-trial interval (ITI, 8-12s) featured a fixation cross. Participants assigned affective ratings for stimuli ‘X’, ‘C’, ‘D’, and the fixation cross immediately following the test phase, providing measures of subjective positive affect and anxious arousal.

The experiment featured conditioning and test phases. Six CS configurations were presented in conditioning: A+, AX-, BC-, D+, AD+, and X- (Figure 1). ‘AX-’ was the conditioned inhibitor, which was compared to the control safety signal ‘BC-’: hereafter referred to as the *standard safety signal*. Each CS was presented for 6 trials each, in pseudorandomized order, resulting in 36 trials. The test phase consisted of three CS configurations (AD+, DC-, DX-) also for 6 trials each, resulting in 18 trials total. To ensure that test responses were not the product of conscious instruction (see (Mechias et al., 2010), the test phase commenced immediately after conditioning with no signaled interlude. At test, the inhibitor (X) and standard safety signal (C) were combined with the threat signal D+ and not reinforced (DX-, DC-), while AD+ was reinforced throughout. To control for the influence of presentation order, approximately half of participants were presented with DX-on their first test trial, with the others presented with DC- (Laing et al., 2021). Following the task’s final phase, participants rated the degree to which the X, C, D, and the fixation cross stimuli, respectively, evoked changes in positive-negative affect and anxious arousal via standardized self-assessment manikins (Bradley and Lang, 1994). The task was programmed in E-Prime (Psychology Software Tools, Pittsburgh, PA) and was presented on a 32” LCD BOLD screen (Cambridge Research Systems) visible via a reverse mirror mounted on the head coil. Noise bursts (US) were delivered via Sensimetrics Insert Earphones (S15 model, Sensimetrics Corp.), which also provided passive noise cancellation (∼30dB). Participants’ responses were registered with a 2-button LS-PAIR Lumina response pad (Cedrus Corporation USA), which they were familiarized with prior to scanning.

### Skin conductance responses (SCR)

Skin conductance was recorded using MRI-compatible finger electrodes (Ag/AgCI) fitted with conductance gel (0.5% saline) to the intermediate phalanges of the index and middle finger of participants’ left hands. Fingers were cleaned with alcohol wipes prior to the attachment of electrodes. The signal was amplified and sampled at 1000 Hz using PowerLab v8.0 (ADInstruments, Dunedin, NZ) and recording was triggered concurrently with the beginning of the experiment and the functional imaging sequence. The Psychophysiological Modelling Toolbox (PsPM, Bach et al., 2018; Bach and Melinscak, 2020) in MATLAB (version 9.4, The MathWorks Inc.) was used for pre-processing and modelling of SCRs. A custom program was used to identify artefacts in the skin conductance timeseries, which were removed in PsPM prior to further analysis. After artefact removal, SC data were filtered with a 10ms median filter followed by a first order bidirectional band-pass Butterworth filer (cut-off frequencies 0.0159 - 5Hz) and down-sampled to 10Hz. Dynamic Causal Modelling was implemented via the PsPM toolbox to provide trial-by-trial estimates of sympathetic nervous system activity. SCR artefacts were detected via a semi-automated process. A custom program identified time series containing: a) signal increases greater than 20% per second, b) signal decreases greater than 10% per second, or c) absolute changes greater than 0.075 µS per millisecond. 17 subjects’ SC data were flagged for review based on these criteria and excluded after manual review. A further 7 subjects’ SCR data was comprised by technical issues. This resulted in a final SCR sample of *N*=24.

### Image acquisition

Imaging was performed on a 7T research scanner (Siemens Healthcare, Erlangen, Germany) equipped with a 32-channel head-coil (Nova Medical Inc., Wilmington MA, USA). The functional sequence consisted of a multi-band (6 times) and grappa (2 times) accelerated GE-EPI sequence in the steady state (TR, 800ms; TE, 22.2 ms; pulse/flip angle, 45°; field-of-view, 20.8 cm; slice thickness (no gap), 1.6 mm; 130 × 130-pixel matrix; 84 interleaved axial slices aligned to AC-PC line; Setsompop et al., 2012). The total sequence time was 16 minutes and 10 seconds, corresponding to 1202 whole-brain EPI volumes. A T1-weighted high-resolution anatomical image (MP2RAGE; Marques et al. 2010) was acquired for each participant to assist with functional time series co-registration (TR = 5000 ms; TE, 3.0 ms; inversion times, 700/2700ms; pulse/flip angles, 4/5°; field-of-view, 24 cm; slice thickness (no gap), 0.73 mm; 330 × 330–pixel matrix; 84 sagittal slices aligned parallel to the midline). The total sequence time was 7 minutes and 12 seconds. To assist with head immobility, foam-padding inserts were placed on either side of the participants’ head. Cardiac and respiratory recordings were sampled at 50 Hertz (Hz) using a Siemens (Bluetooth) pulse-oximeter and respiratory belt. Information derived from these recordings were used for physiological noise correction (see further).

### Image pre-processing

Imaging data was pre-processed using Statistical Parametric Mapping (SPM) 12 (v7771, Welcome Trust Centre for Neuroimaging, London, UK) within a MATLAB 2019b environment (The MathWorks Inc., Natick, MA). Motion correction was performed by realigning each subject’s time series to the mean image, and all images were resampled using 4th Degree B-Spline interpolation. Individualized motion parameters were estimated via Motion Fingerprint (Wilke, 2012) to account for head motion, and participant data was excluded if mean total scan-to-scan voxel displacement exceeded 1.6 mm (one voxel), resulting in the exclusion of four participants. Each participant’s anatomical image was co-registered to their respective mean functional image, segmented, and normalized to the International Consortium of Brain Mapping template using the unified segmentation plus DARTEL approach. Smoothing was applied with a 3.2mm^3^ full-width-at-half-maximum (FWHM) gaussian kernel to increase anatomical precision. Physiological noise was modelled at the first level using the PhysIO Toolbox (Kasper et al., 2017). This toolbox applies noise correction to fMRI sequences using physiological recordings and is shown to enhance blood-oxygen level dependent (BOLD) signal sensitivity and temporal signal-to-noise ratio (tSNR) at 7T (Reynaud et al., 2017). The Retrospective Image-based Correction function (RETROICOR, Glover et al., 2000) was applied to model periodic effects of heartbeat and respiration on BOLD signals. The Respiratory Response Function (RRF, Birn et al., 2008), convolved with respiration volume per time (RVT), was used to model low frequency signal fluctuations arising from changes in depth and rate of breath. Heart rate variability (HRV) was convolved with a Cardiac Response Function (CRF, Chang et al., 2009) to account for BOLD variances due to heartrate-dependent changes in blood oxygenation. Individualized DARTEL tissue maps segmented from each participant’s respective anatomical scan were used to apply aCompCor, which models negative BOLD signals using principal components derived from white matter and CSF (Behzadi et al., 2007).

### fMRI analyses

Each participant’s pre-processed time-series was included in a first-level SPM general linear model (GLM) analysis, which specified the onsets of each CS event-type (grouped as separate conditions for ‘early’/first 3 trials and ‘late’/last 3 trials) in each task phase to be convolved with canonical hemodynamic response function. The fixation-cross ITI periods throughout whole task served as the implicit baseline. A high-pass filter (1/128 s) accounted for low-frequency noise, while temporal autocorrelations were estimated with an autoregressive model. Contrast images were estimated for each CS condition against the implicit baseline and were carried forward to the group-level using the summary statistics approach to random-effects analyses.

As our first aim was to characterize differences in brain response to inhibitory versus standard safety signals, we compared all trials of the conditioned inhibitor and standard safety signal during the conditioning phase (AX- vs. BC-). Secondly, to identify learning-specific responses, we compared early versus late trials for each safety signal, respectively (e.g., AX- _*early*_ vs. AX- _*late*_). Thirdly, to distinguish brain responses to each safety signal during learning versus expression, we compared responses to each stimulus type across conditioning and test phases (e.g., AX- vs. DX-), and differences between safety signals when combined with a CS+ at test (e.g., DX- vs. DC-). For each of the main conditioned inhibitor contrasts, we provide extended results that illustrate its overlap with brain responses to the simple conditioned threat (A+). Statistical significance was estimated with a whole-brain false-discovery rate corrected threshold (*P*_FDR_ < 0.05) and a 5-voxel cluster-extent threshold (K_E_ ≥ 5). Analyses of threat-expectancy ratings and SCR were performed by separating responses into early and late phases (3 trials each) consistent with the imaging analyses, and subjected to repeated-measures ANOVAs and post-hoc t-tests. One-sample t-tests were used to compare differences in subjective ratings (valence and arousal) that were collected at the end of the task.

## Results

### BEHAVIORAL RESULTS

#### Safety learning: conditioned inhibitor versus standard safety signal

Conditioned response measures (threat-expectancy ratings and SCR) were analyzed to assess changes in threat response across the conditioning phase. Threat-expectancy ratings for the conditioned inhibitor and control cue showed a main effect of stimulus (*F*_1, 45_ = 35.22, *p* < 0.001, *η*^2^_*p*_ = 0.44) and phase (early, late, *F*_1, 45_ = 234.45, *p* < 0.001, *η*^2^_*p*_ = 0.84), but no interaction. Post-hoc test showed higher threat-expectancy for the inhibitor (AX- > BC-: *M* = 10, *t* = 5.93, *d* = 0.87, *p* < 0.001), driven by significant differences in early (*M* = 14.85, *t* = 6.47, *p* < 0.001), but not late (*M* = 5.15, *t* = 2.24, *p* = 0.17) conditioning trials (Figure 2A). SCRs (Figure 2B) showed no effect of stimulus (AX- > BC-; *p* = 0.93), but a significant effect of phase (early, late, *F*_1, 23_ = 4.58, *p* < 0.05, *η*^2^_*p*_ = 0.17), indicating elevated SCRs in early conditioning which were decreased by later trials (early > late: *M* = 0.13, *t* = 2.14, *d* = 0.44, *p* = 0.04). Transfer test responses showed mixed results. Threat-expectancy showed main effects of stimulus (*F*_1, 45_ = 894.58, *p* < 0.001, *η*^2^_*p*_ = 0.95), phase (*F*_1, 45_ = 28.25, *p* < 0.001, *η*^2^_*p*_ = 0.39), and their interaction (*F*_1, 45_ = 16.48, *p* < 0.001, *η*^2^_*p*_ = 0.27). Expectancy was robustly decreased for the conditioned inhibitor (AD+ > DX-; *M* = 88.08, *t* = 36.74, *d* = 5.42, *p* < 0.001) and standard safety signal (AD+ > DC-; *M* = 87.57, *t* = 36.52, *d* = 5.38, *p* < 0.001) relative to the threat compound, but showed no significant differences between them (DX- vs. DC-). In SCRs, no main effects were identified for stimuli (AD+, DX-, DC-; *p* = 0.27) or phase (early, late, *p* = 0.96). In sum, behavioral measures indicated that each safety signal evoked equivalent decreases in behavioural threat response during learning, but did not differ in responses evoked at test.

**Figure 2.**
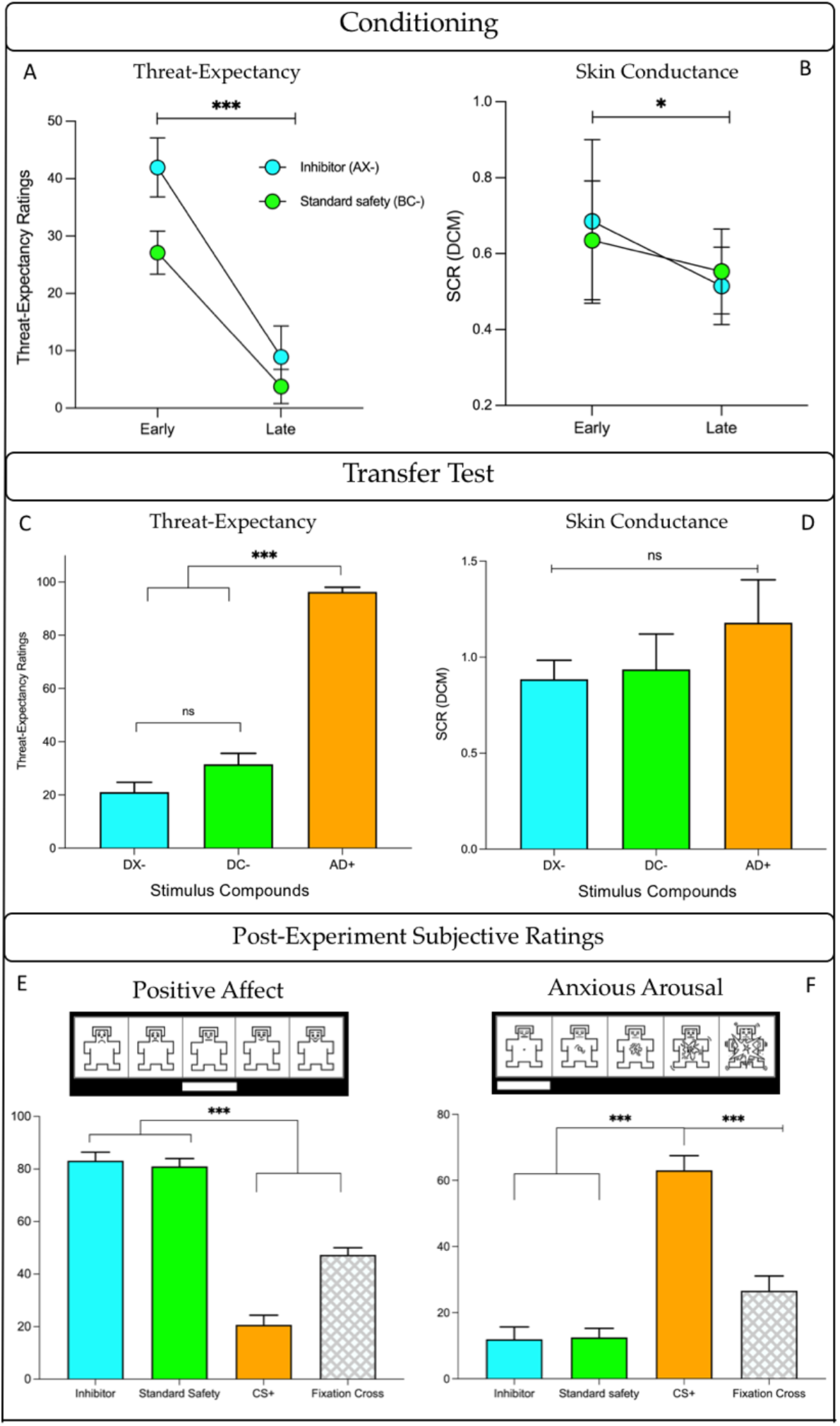
Behavioral measures. (A) Threat-expectancy ratings showed a significant decreased across conditioning for each safety signal, with higher ratings during early trials for the inhibitor versus standard, and no difference during late conditioning. (B) SCRs showed no difference between stimuli (inhibitor vs. standard), but a significant decrease from early to late conditioning for each stimulus. (C) Both safety signals inhibited threat expectancy at test compared to the threat compound AD+, but did not differ from one another. (D) SCRs showed no differences at test. (E) Subjective affective ratings reliably discriminated safety signals with high positive affect from the CS+ and neutral fixation cross. (F) Similarly, safety signals elicited low appraisal of anxious arousal relative to high ratings of CS+, and moderate ratings of the fixation cross. ‘AX-’ = conditioned inhibitor. ‘BC-’ = standard safety signal. ‘DX-’ = inhibitor + threat cue. ‘DC’ = standard safety signal + threat cue. ‘AD+’ = threat compound. ‘DCM’ = dynamic causal model for SCR (via PsPM). Error bars represent 95% confidence intervals. * = *p* < 0.05, *** = *p* < 0.001, ns = *p* > 0.05.

#### Subjective ratings: positive affect and anxious arousal

Compared to a CS+, the conditioned inhibitor and standard safety signal elicited greater positive affect (X > D: *M* = 62.5, t = 7.58, *d* = 1.12, *p* < 0.001; C > D: *M* = 60.33, *t* = 7.32, *d* = 1.08, *p* < 0.001; Figure 2E) and lower anxious arousal (X > D: *M* = 51.09, *t* = 10.38, *d* = 1.53, *p* < 0.001; C > D: *M* = 50.54, *t* = 10.26, *d* = 1.51, *p* < 0.001; Figure 2F). Further, both safety signals also evoked greater positive affect (X > fixation cross: *M* = 26.09, *t* = 3.17, *d* = 0.47, *p* = 0.006; C > fixation cross: *M* = 23.92, *t* = 2.90, *d* = 0.43, *p* = 0.009; Figure 2E) and lower arousal (fixation cross > X: *M* = 14.67, *t* = 2.98, *d* = 0.44, *p* = 0.01; fixation cross > C: *M* = 14.13, *t* = 2.87, *d* = 0.42, *p* = 0.01; Figure 2F) relative to ratings of the fixation cross, which served as a putative baseline comparison. The inhibitor and standard safety signal did not differ from one another on either affective measure. In summary, both safety signals accumulated high positive affect and low anxious arousal following conditioning, consistent with previous findings (Harrison et al., 2017; Laing et al., 2021).

### IMAGING RESULTS

#### Differential safety responses: conditioned inhibitor versus standard safety signal

We first estimated differential neural responses to the conditioned inhibitor versus standard safety signal (AX- > BC-). As shown in Figure 3, this direct comparison identified significantly greater activation to the conditioned inhibitor in regions including the anterior insular cortex bilaterally, extending to the dorsal putamen in the right hemisphere; the left caudate body extending to dorsal putamen, the left ventrolateral cerebellum (Crus II), and right posterior dorsolateral prefrontal cortex (Figure 3). Figure 3-1 (extended data) highlights the overlap between the differential response to the conditioned inhibitor versus standard safety signal and the response to conditioned threat alone (A+). Partial overlap was noted with regards to activation of the anterior insular cortex. The standard safety signal compared to conditioned inhibitor (BC- > AX-) evoked significantly greater activation of the ventromedial prefrontal cortex, spanning posterior (subgenual) and anterior (frontopolar) subregions, retrosplenial posterior cingulate cortex, right posterior hippocampus, basal forebrain (∼medial forebrain bundle), right posterior primary motor cortex, cerebellum (VI and VIIb) medial and lateral visual association cortex, spanning the fusiform, lingual gyrus and occipital pole. Complete results are provided in Table 1. All GLM results are presented on the ‘Synthesized_FLASH25’ (500um, MNI space) *ex vivo* template (Edlow et al., 2019).

**Table 1.**
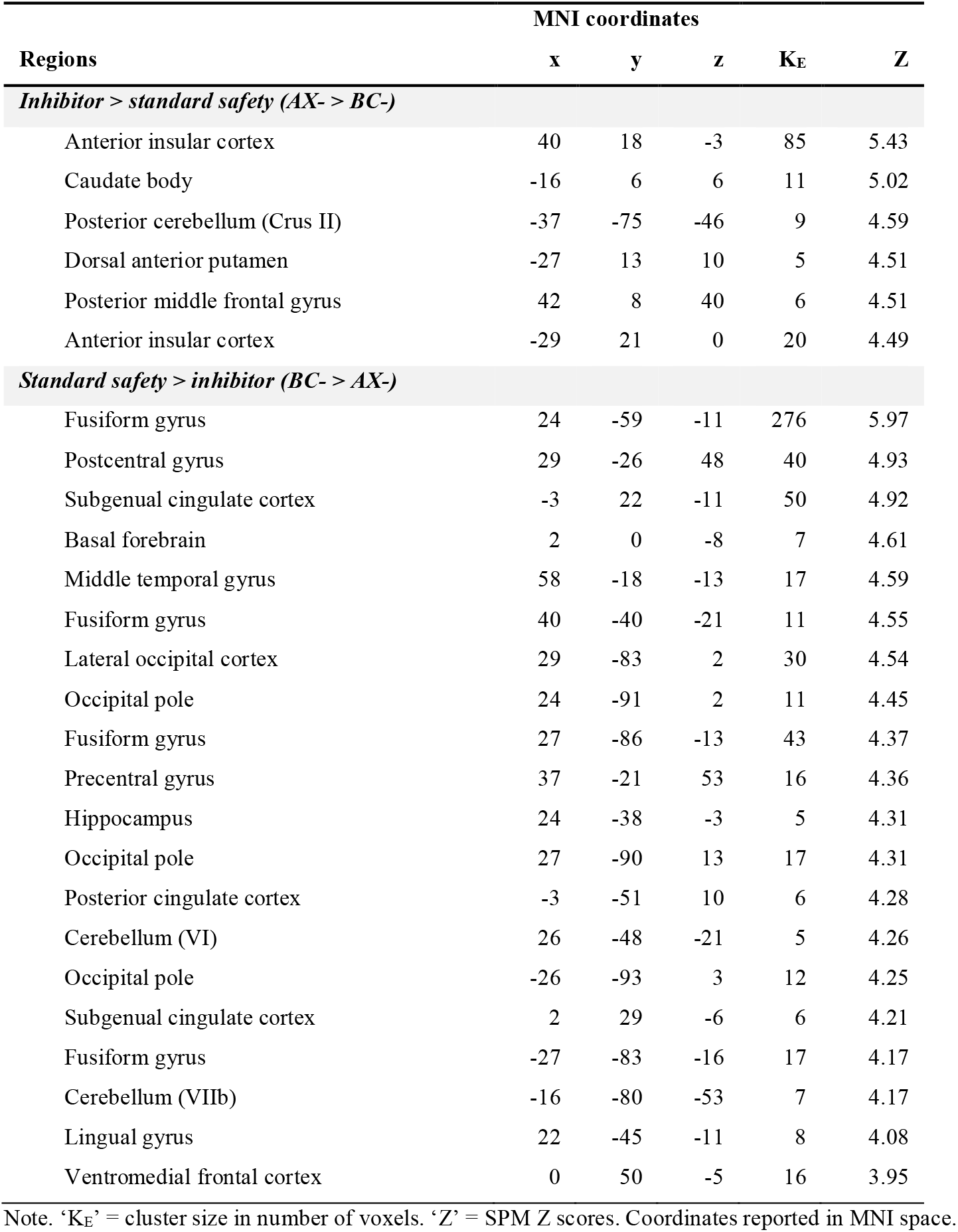
Differential safety responses – conditioned inhibitor versus standard safety signal. Whole brain contrasts estimated at *P*_FDR <_ 0.05.

| Regions | MNI coordinates |  |  |  |  |
| --- | --- | --- | --- | --- | --- |
|  | x | y | z | K<sub>E</sub> | Z |
| *Inhibitor > standard safety (AX- > BC-)* |  |  |  |  |  |
| Anterior insular cortex | 40 | 18 | -3 | 85 | 5.43 |
| Caudate body | -16 | 6 | 6 | 11 | 5.02 |
| Posterior cerebellum (Crus II) | -37 | -75 | -46 | 9 | 4.59 |
| Dorsal anterior putamen | -27 | 13 | 10 | 5 | 4.51 |
| Posterior middle frontal gyrus | 42 | 8 | 40 | 6 | 4.51 |
| Anterior insular cortex | -29 | 21 | 0 | 20 | 4.49 |
| *Standard safety > inhibitor (BC- > AX-)* |  |  |  |  |  |
| Fusiform gyrus | 24 | -59 | -11 | 276 | 5.97 |
| Postcentral gyrus | 29 | -26 | 48 | 40 | 4.93 |
| Subgenual cingulate cortex | -3 | 22 | -11 | 50 | 4.92 |
| Basal forebrain | 2 | 0 | -8 | 7 | 4.61 |
| Middle temporal gyrus | 58 | -18 | -13 | 17 | 4.59 |
| Fusiform gyrus | 40 | -40 | -21 | 11 | 4.55 |
| Lateral occipital cortex | 29 | -83 | 2 | 30 | 4.54 |
| Occipital pole | 24 | -91 | 2 | 11 | 4.45 |
| Fusiform gyrus | 27 | -86 | -13 | 43 | 4.37 |
| Precentral gyrus | 37 | -21 | 53 | 16 | 4.36 |
| Hippocampus | 24 | -38 | -3 | 5 | 4.31 |
| Occipital pole | 27 | -90 | 13 | 17 | 4.31 |
| Posterior cingulate cortex | -3 | -51 | 10 | 6 | 4.28 |
| Cerebellum (VI) | 26 | -48 | -21 | 5 | 4.26 |
| Occipital pole | -26 | -93 | 3 | 12 | 4.25 |
| Subgenual cingulate cortex | 2 | 29 | -6 | 6 | 4.21 |
| Fusiform gyrus | -27 | -83 | -16 | 17 | 4.17 |
| Cerebellum (VIIb) | -16 | -80 | -53 | 7 | 4.17 |
| Lingual gyrus | 22 | -45 | -11 | 8 | 4.08 |
| Ventromedial frontal cortex | 0 | 50 | -5 | 16 | 3.95 |

Note. ‘ $K_E$ ’ = cluster size in number of voxels. ‘ $Z$ ’ = SPM $Z$ scores. Coordinates reported in MNI space.

**Figure 3.**
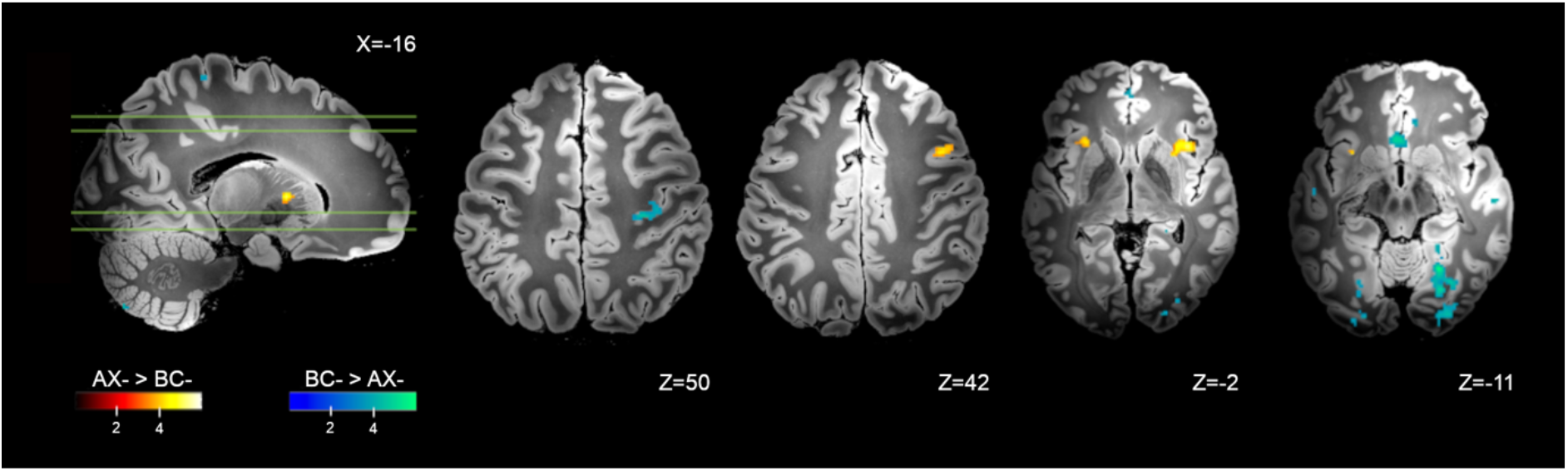
Brain regions with significant differential response to the conditioned inhibitor (AX- > BC-) versus the standard safety signal (BC- > AX-). Whole-brain FDR-corrected (*P* < 0.05) results are displayed on a high-resolution anatomical template in MNI space.

#### Safety learning dynamics during conditioning

Safety learning dynamics were analyzed via contrasts of early versus late trials *within* each safety stimulus during conditioning. As shown in Figure 4, early versus late trials for the conditioned inhibitor (AX- _*early*_ > AX- _*late*_), were associated with significantly greater activation of the caudate body extending to dorsal anterior putamen and globus pallidus (external), the midbrain substantia nigra, ventral lateral and intralaminar thalamic nuclei, dorsal-mid cingulate and dorsal premotor cortex, and the cerebellum (IX). There was no significantly greater activation identified for the conditioned inhibitor during late vs early trials (AX- _*late*_ > AX- _*early*_). Figure 4-1 (extended data) illustrates that there was minimal overlap between the differential response to the early versus late conditioned inhibitor trials and the response to the conditioned threat alone (A+).

**Figure 4.**
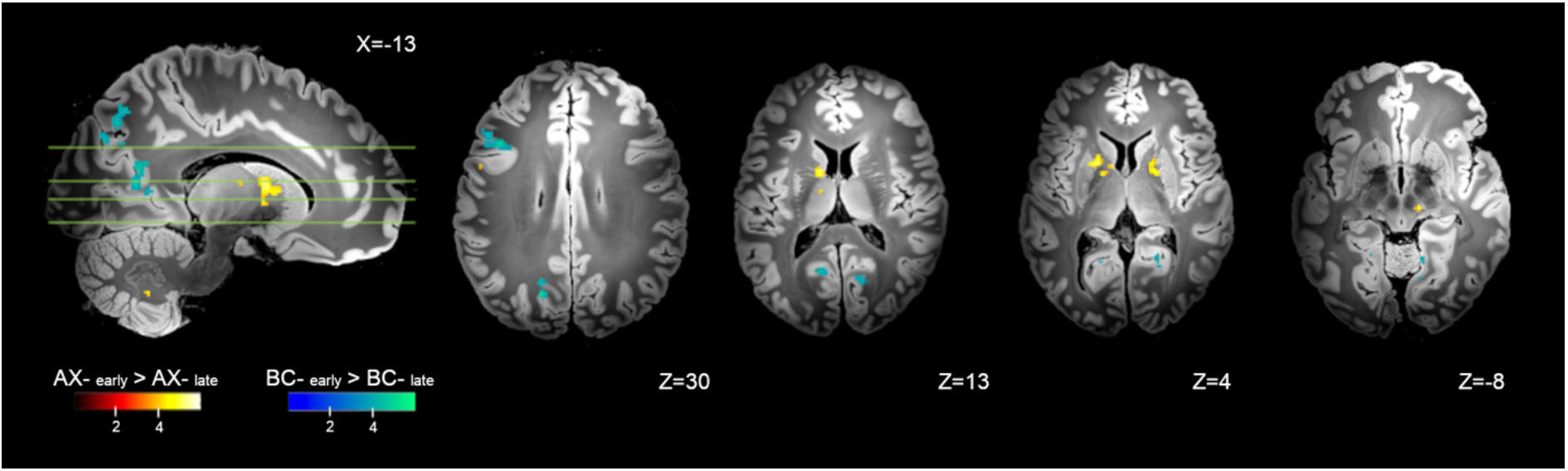
Brain regions with significant differential response to early versus late conditioning trials, for the conditioned inhibitor (AX- _*early*_ > AX- _*late*_) and standard safety signal (BC- _*early*_ > BC- _*late*_). Whole-brain FDR-corrected (*P* < 0.05) results are displayed on a high-resolution anatomical template in MNI space.

For the standard safety signal (BC- _*early*_ > BC- _*late*_), early trials evoked significantly greater activation of posterior dorsolateral prefrontal cortex, superior- and intraparietal cortex, precuneus (dorsal and ventral subareas), and medial and lateral visual association cortex, spanning the fusiform, lingual gyrus and occipital pole. There was no significantly greater activation identified for the standard safety signal during late vs early trials (BC- _*late*_ > BC- _*early*_). Complete results are provided in Table 2.

**Table 2.** Safety learning dynamics – early versus late conditioning. Whole brain contrasts estimated at *P*_FDR <_ 0.05.

| Regions | MNI coordinates | | | $K_E$ | $Z$ |
| --- | --- | --- | --- | --- | --- |
|  | x | y | z |  |  |
| *Conditioned inhibitor (AX- early > AX- late)* |  |  |  |  |  |
| Caudate body | -16 | 6 | 10 | 213 | 5.68 |
| Midbrain (substantia nigra) | 11 | -22 | -8 | 10 | 5.13 |
| Pallidum | 19 | 2 | 3 | 128 | 4.90 |
| Thalamus (intralaminar) | 8 | -22 | -2 | 9 | 4.61 |
| Thalamus (ventral lateral nucleus) | -11 | -10 | 13 | 10 | 4.44 |
| Cerebellum (IX) | -13 | -56 | -43 | 7 | 4.35 |
| Mid cingulate cortex | -6 | 19 | 43 | 8 | 4.30 |
| Precentral gyrus | -53 | 5 | 29 | 9 | 4.30 |
| *Standard safety (BC- early > BC- late)* |  |  |  |  |  |
| Superior frontal gyrus | -21 | 8 | 51 | 78 | 5.12 |
| Superior parietal lobule | -10 | -69 | 50 | 179 | 5.10 |
| Lingual gyrus | 19 | -45 | -11 | 31 | 5.06 |
| Precuneus | -18 | -70 | 30 | 33 | 5.03 |
| Ventral precuneus | -11 | -62 | 11 | 93 | 4.87 |
| Middle frontal gyrus | 38 | 22 | 26 | 30 | 4.81 |
| Middle frontal gyrus area | -42 | 18 | 30 | 120 | 4.78 |
| Lingual gyrus | 14 | -67 | -5 | 49 | 4.75 |
| Precuneus | -18 | -66 | 22 | 34 | 4.71 |
| Ventral precuneus | 21 | -56 | 8 | 62 | 4.65 |
| Precuneus | -18 | -64 | 29 | 14 | 4.57 |
| Superior parietal lobule | 11 | -69 | 48 | 33 | 4.55 |
| Ventral precuneus | -16 | -58 | 6 | 25 | 4.44 |
| Supracalcarine cortex | 6 | -77 | 18 | 17 | 4.37 |
| Intracalcarine sulcus | 16 | -69 | 11 | 75 | 4.36 |
| Lateral occipital cortex | 19 | -82 | 27 | 12 | 4.24 |
| Superior parietal lobule | 13 | -46 | 58 | 11 | 4.18 |
| Fusiform gyrus | -24 | -46 | -13 | 7 | 4.15 |
| Lingual gyrus | 13 | -56 | -8 | 10 | 4.13 |
| Fusiform gyrus | -30 | -42 | -16 | 5 | 4.12 |
| Occipital pole | -11 | -91 | 26 | 6 | 4.10 |
| Precuneus | 16 | -64 | 38 | 9 | 4.06 |
| Precuneus | 16 | -66 | 22 | 7 | 4.02 |
| Cuneus | 13 | -85 | 29 | 6 | 4.00 |
| Precuneus | 11 | -67 | 27 | 13 | 3.96 |
| Lingual gyrus | 24 | -51 | -10 | 5 | 3.92 |
| Lingual gyrus | -19 | -54 | -11 | 6 | 3.88 |
| Intraparietal sulcus | -21 | -75 | 43 | 7 | 3.78 |

Note. ' $K_E$ ' = cluster size in number of voxels. ' $Z$ ' = SPM Z scores. Coordinates reported in MNI space.

#### Conditioning versus test

As shown in Figure 5, direct comparison of the conditioned inhibitor across the conditioning and test phases (AX- > DX-) identified significantly greater activation of the bilateral dorsal anterior putamen and globus pallidus (external), the left caudate (tail), midbrain substantia nigra and periaqueductal grey, bilateral dorsal premotor cortex, and precuneus. Complete results are provided in Table 3. Figure 5-1 (extended data) highlights the overlap between the differential response to the conditioned inhibitor at training versus test and the response to the conditioned threat alone (A+). Partial overlap was noted with regards to activation of the left premotor cortex. There was no significantly greater activation identified for the conditioned inhibitor at test compared to conditioning (DX- > AX-). For the standard safety signal, significantly greater activation of the anterior fusiform and lingual gyrus was identified during conditioning compared to test (BC- > DC-). There was no significantly greater activation identified for the standard safety signal at test compared to conditioning (DC- > BC-). As a further test of safety expression, we contrasted test-phase responses to the inhibitor and standard safety signal directly (DX- vs. DC-). No clusters exceeded the *P*_FDR <_ 0.05 threshold (K_E_ ≥ 5 voxels).

**Table 3.** Safety conditioning versus test phase. Whole brain contrasts estimated at *P*_FDR <_ 0.05.

| Regions | MNI coordinates |  |  |  |  |
| --- | --- | --- | --- | --- | --- |
|  | x | y | z | K<sub>E</sub> | Z |
| *Conditioned inhibitor (AX- > DX-)* |  |  |  |  |  |
| Putamen | -14 | 6 | 6 | 371 | 6.07 |
| Putamen | 26 | 2 | 0 | 269 | 5.54 |
| Midbrain (substantia nigra) | 18 | -16 | -13 | 19 | 5.54 |
| Superior frontal gyrus | -19 | 3 | 51 | 138 | 4.97 |
| Middle frontal gyrus | 26 | 27 | 32 | 19 | 4.71 |
| Brainstem (periaqueductal grey) | 8 | -35 | -6 | 10 | 4.57 |
| Superior frontal gyrus | -26 | 2 | 46 | 14 | 4.46 |
| Caudate body | 18 | 10 | 10 | 20 | 4.38 |
| Precuneus | 19 | -54 | 16 | 8 | 4.34 |
| Caudate tail | -19 | -2 | 18 | 5 | 4.27 |
| Superior frontal gyrus | -21 | 21 | 51 | 7 | 4.02 |
| Middle frontal gyrus | 30 | 37 | 29 | 7 | 3.98 |
| Superior frontal gyrus | 24 | 11 | 56 | 6 | 3.89 |
| *Standard safety (BC- > DC-)* |  |  |  |  |  |
| Precentral gyrus | 40 | -13 | 56 | 41 | 5.93 |
| Fusiform gyrus | 22 | -45 | -13 | 48 | 5.48 |
| Lingual gyrus | 11 | -62 | 2 | 9 | 4.65 |
| Lingual gyrus | 22 | -59 | -10 | 5 | 4.49 |

Note. ' $K_E$ ' = cluster size in number of voxels. ' $Z$ ' = SPM Z scores. Coordinates reported in MNI space.

**Figure 5.**
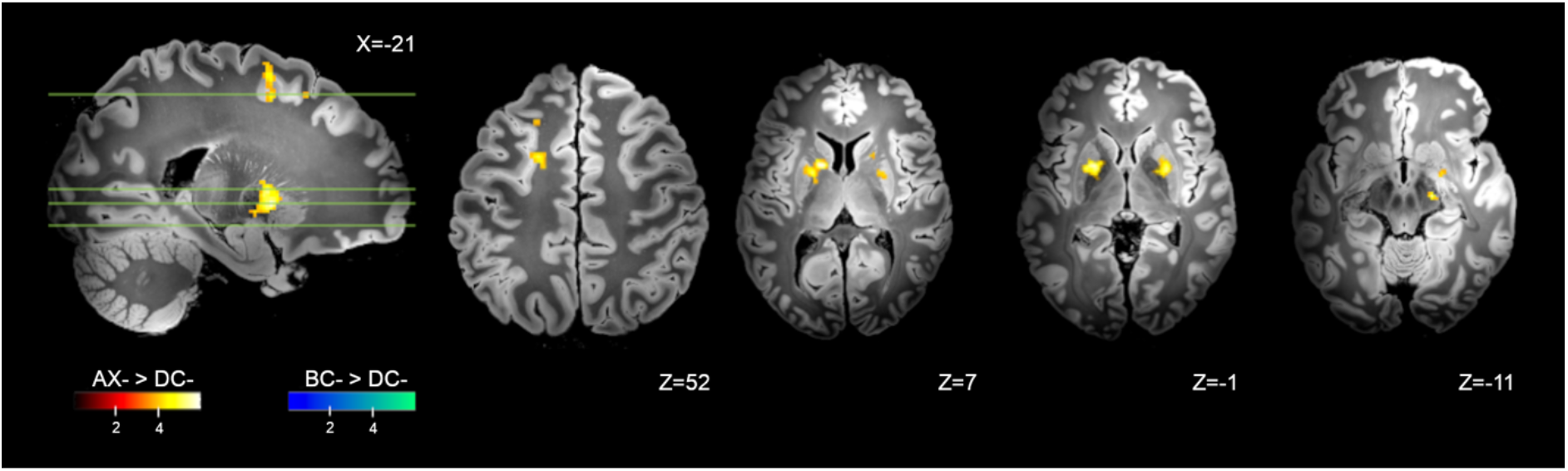
Brain regions with significant differential response during conditioning versus transfer test, for the conditioned inhibitor (AX- > DX-) and standard safety signal (BC- > DC-). Whole-brain FDR-corrected (*P* < 0.05) results are displayed on a high-resolution anatomical template in MNI space.

## Discussion

This study combined ultra-high field fMRI with Pavlovian conditioned inhibition to characterize the neural basis of safety learning in humans. We compared two kinds of conditioned safety signals: a conditioned inhibitor, which preceded threat-omission in compound with a CS+ (A+, AX-), and a standard safety signal, a stimulus compound that was unreinforced (BC-) without threat-proximity. Supporting our general hypotheses, safety learning via conditioned inhibition evoked prominent subcortical activations spanning the dorsal striatum, thalamus, and midbrain, together with dorsal prefrontal and premotor cortex regions. Conversely, standard safety signal processing was associated with greater engagement of distributed frontal and occipital-parietal regions, which have previously been linked to the cognitive appraisal of safety information.

Compared to the standard safety signal, the conditioned inhibitor selectively engaged the dorsal anterior striatum, anterior insular and posterior dorsolateral prefrontal cortex, with further analyses implicating extended activation of midbrain, thalamic and premotor cortex subregions during early safety learning and when comparing conditioning to test phase responses. We observed that activation of the anterior insular, premotor, and midbrain subregions by the conditioned inhibitor overlapped with responses to the conditioned threat signal (A+), whereas most subcortical regions, particularly the dorsal striatum, were non-overlapping. Consequently, the conditioned inhibitor’s neural response encompassed both threat-responsive regions and learning-related cortico-striatal activation. The former may reflect a key component of inhibitory learning – the direct conflict between threat and safety information. Conditioned inhibition depends on a reasonable level of threat expectation (Harris et al., 2014; Lysle and Fowler, 1985), requiring that inhibitory stimuli predict safety when threat is otherwise expected (Sosa and Ramírez, 2019). Insular activity may reflect modulation of threat value (Sharvit et al., 2018; Teckentrup et al., 2019), analogous to its role in fear extinction (Fullana et al., 2018). Interestingly, the non-overlapping cortico-striatal regions associated with conditioned inhibition are constituents of well-established cortico-basal ganglia pathways, which hold deep intrinsic connectivity in primate neuroanatomy. For instance, the dorsal striatum receives inputs from premotor cortical regions, which relays information to the ventrolateral thalamus by way of the pallidum and SNc (Alexander and Crutcher, 1990; Alexander et al., 1986; Haber and Knutson, 2010). Anatomical characteristics of this circuit are also consistent with intrinsic premotor cortex connectivity with the dorsal striatum and SNc observed in humans via fMRI (Choi et al., 2012; Di Martino et al., 2008; Morris et al., 2016).

The engagement of these cortico-striatal regions by the inhibitor, but not the standard safety signal, is consistent with the primary psychological difference between these stimuli: the manipulation of expectancy violation (Laing et al., 2021). In our task, early trials of the conditioned inhibitor (AX-) are expected to evoke prediction error-like mismatches, wherein threat-anticipation elicited by ‘A+’ is violated by threat-omission following ‘AX-’ (Laing and Harrison, 2021). These trials demand the reconciliation of expected (threat) and actual (safety) outcomes, but as the stimulus-safety contingency is repeated, the need for error-processing and value-updating is minimized. Consequently, the observed temporal shift in dorsal striatal activity, in tandem with SNc, is consistent with the role of prediction error in associative learning theory (Pauli et al., 2015; Schultz and Dickinson, 2000; Schultz et al., 2003). For instance, dorsal striatal nuclei show elevated prediction error responses in early conditioning, which decreases as contingencies are learned (Cooper et al., 2012; Oyama et al., 2015; Valentin and O’Doherty, 2009), while also updating conditioned responses to the CS (Yoshizawa et al., 2018). The dorsal striatum receives direct projections from midbrain dopaminergic neurons of the SNc (Lanciego et al., 2012), which provide a prediction error signal by firing in response to unexpected stimulus-delivery (or omission), and decreasing in magnitude as expectations and outcomes are aligned (Eshel et al., 2016; Salinas-Hernández et al., 2018; Waelti et al., 2001). As a result, safety learning via conditioned inhibition appears to invoke a more direct instantiation of these active error-corrective learning systems compared to the standard safety signal, which is passively unreinforced, and does not directly incur an expectancy-outcome mismatch. Our findings thereby implicate a specific role for these associative learning mechanisms in the domain of human safety learning, which has been primarily associated with prefrontal cortical mechanisms (Tashjian et al., 2021).

Brain regions associated with the standard safety signal (BC-) overlap with the results of differential fear conditioning studies, which report consistent involvement of the vmPFC, hippocampus, and posterior cingulate cortex in safety versus threat discrimination (Fullana et al., 2015). However, as noted earlier, these experiments were not designed to discriminate mechanisms of safety learning from safety expression and have likely conflated these processes. An acquisition-expression dissociation is well-supported in animal models, which demonstrate learning-specific roles for the striatum and insular (Foilb et al., 2016; Rogan et al., 2005), and expression-specific contributions of hippocampal and prefrontal cortical regions (Kreutzmann and Fendt, 2020; Kreutzmann et al., 2020; Meyer et al., 2019; Yan et al., 2019). In humans, learned safety should facilitate experiences of positive affect and inhibition of fear behaviors (Zhang et al., 2015), each requiring recall of safety information from memory. Affective valuation, response-inhibition, and recall processes all converge on these standard safety-processing regions (Harrison et al., 2017; Hennings et al., 2020; Hermann et al., 2020; Merz et al., 2018; Roy et al., 2012). The vmPFC has a notably multifaceted functional role, modulating both fear and safety expression, rather than unidirectional threat-inhibition (Battaglia et al., 2021; Tashjian et al., 2021). For instance, it is causally implicated in acquiring conditioned responses to a CS+ (Battaglia et al., 2020), and in inhibiting responses in the face of *unlearned* safety signals, (e.g., familial attachment figures) (Eisenberger et al., 2011). Thus, a ‘dual systems’ perspective on human safety learning may be warranted, featuring (i) a cortico-striatal system that encodes initial safety learning associations, and (ii) an expanded cortical system supporting the appraisal and expression (or ‘use’) of learned safety information. Synchronization between these systems likely requires interactions extending beyond conventional vmPFC-oriented circuitry, involving midbrain, striatal, and prefrontal integration of both prediction error-based learning and safety-retention across time (Esser et al., 2021; Gerlicher et al., 2018; Raczka et al., 2011; Thiele et al., 2021; Yan et al., 2019).

Although few, if any, existing fMRI studies have translated the Pavlovian conditioned inhibition model to human fear conditioning, it has recently been reported in a study of appetitive conditioning, where the inhibitor predicted reward-omission, and evoked dorsal-striatal responses similar to those observed here (Mollick et al., 2021). Consistent with associative learning theory (Roesch et al., 2012), these overlapping findings suggest that US-omission can function as a potent reinforcer when it contradicts expectation (Tobler et al., 2003), rather than being a meaningless or neutral event. The convergence of threat and reward-omission on dorsal striatal systems thus points to a more fundamental role in inhibitory learning, such that the putamen, caudate, and pallidum encode learning of ‘CS → no US’ associations whether the omitted-US is pleasant or noxious. In comparison, other studies suggest an opposing role for the ventral *striatum*, which underlies the dynamic reorganization of aversive and rewarding *excitatory* (CS → US) associations (Grill et al., 2021; Klucken et al., 2009; Li et al., 2011; Richter et al., 2020; Schiller et al., 2008; Stelly et al., 2021). Similarly, dopamine signals in the nucleus accumbens do not appear to encode prediction errors for threat-omission (Kutlu et al., 2021), which may explain the ventral striatum’s lack of involvement in safety learning (Josselyn et al., 2005; Mohammadi et al., 2014).

While comparisons between the training and test phases were instructive for discriminating safety learning from expression related neural activity, we did not observe robust behavioral differences between the safety signals during the test phase comparison, unlike our recent behavioral study (Laing et al., 2021). This null finding likely reflects adjustments to the task that were made for the fMRI environment, particularly the reduction of trial repetitions to reduce the overall length of the experiment. It may also be useful to examine test responses in future applications of the conditioned inhibition approach after imposing a delay between the conditioning and test phase (e.g., ≥ 24hr intervals post-learning, Lonsdorf et al., 2017).

## Conclusion

Overall, our mapping of safety learning via conditioned inhibition aligns with evidence from non-human studies, provides novel evidence for the involvement of known reinforcement learning systems, and proposes an extension to current models of human safety learning. A distributed cortico-striatal system, centered on the dorsal striatum, showed elevated activity during early learning, when threat expectations and safety-outcomes require greatest reconciliation, with decreased activity as trials proceeded. This may represent a neural substrate of safety *learning*, where initial ‘stimulus-safety’ associations are formed, which can be distinguished from wider cortical correlates of safety *expression* that facilitate the behavioral outcomes of learning. These cortico-striatal systems could provide novel avenues for clinical translation. For instance, though anxiety-related neuropathology is often described in terms of prefrontal threat-processing (Alexandra Kredlow et al., 2021), recent studies have shifted focus towards reinforcement learning systems similar to those observed here (Ney et al., 2021; Seidemann et al., 2021; Zilcha-Mano et al., 2020), which could provide new insights for characterizing pathophysiology and neural mechanisms of treatment response.

## Acknowledgments

This work was funded by a National Health and Medical Research Council of Australia (NHMRC) Project Grant (1161897) to BJH. We acknowledge the scientific and technical assistance of the Australian National Imaging Facility – a National Collaborative Research Infrastructure Strategy (NCRIS) capability at the Melbourne Brain Centre Imaging Unit (MBCIU), The University of Melbourne. The multiband fMRI sequence was generously supported by a research collaboration agreement with CMRR, The University of Minnesota and the MP2RAGE works in progress sequence was provided by Siemens Healthineers (Germany). The authors gratefully acknowledge Ms Lisa Incerti for her contributions to data collection.

## Extended Data

**Figure 3-1.**
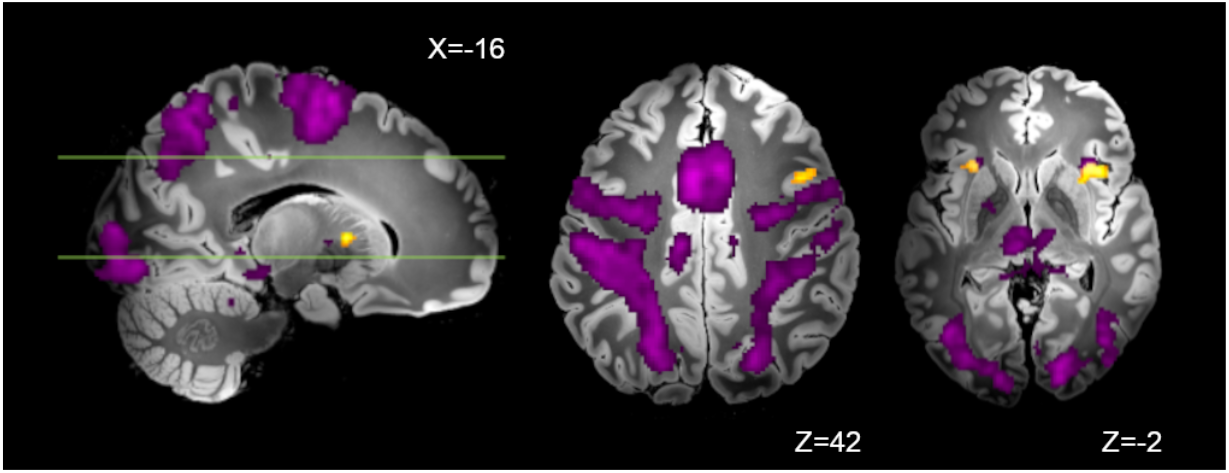
Partial overlap between the differential response to the conditioned inhibitor versus standard safety signal (AX- > BC-, yellow), and the response to the conditioned threat alone (A+, purple) in activation of the anterior insular cortex.

**Figure 4-1.**
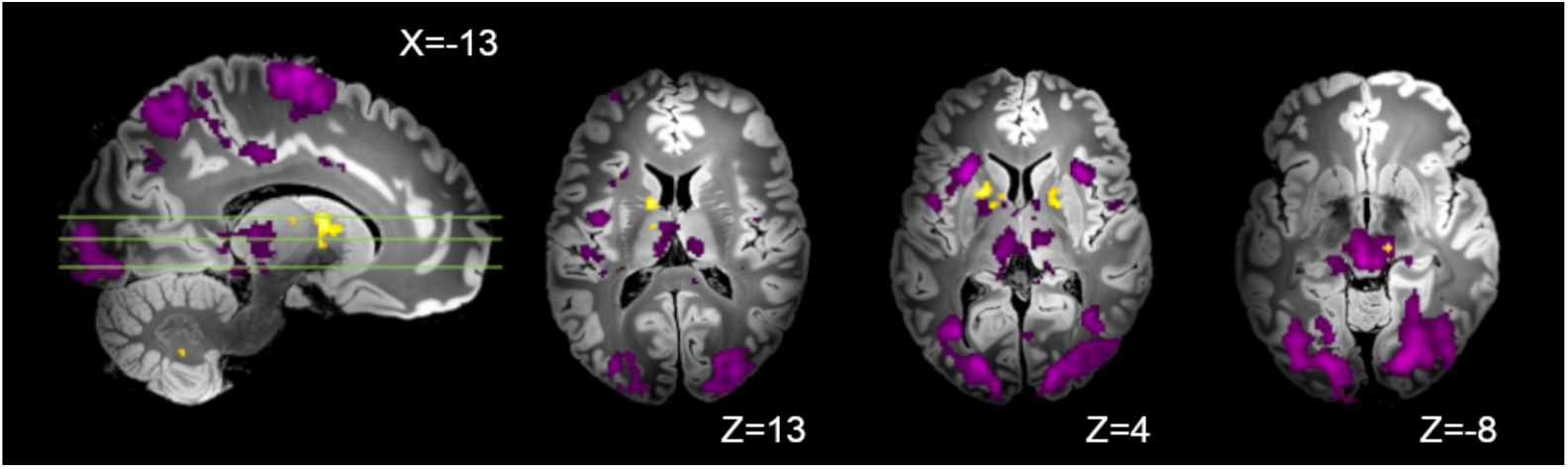
Minimal overlap observed between the differential response to the early versus late conditioned inhibitor trials (AX- _*early*_ > AX-_*late*_, yellow), and the response to the conditioned threat alone (A+, purple).

**Figure 5-1.**
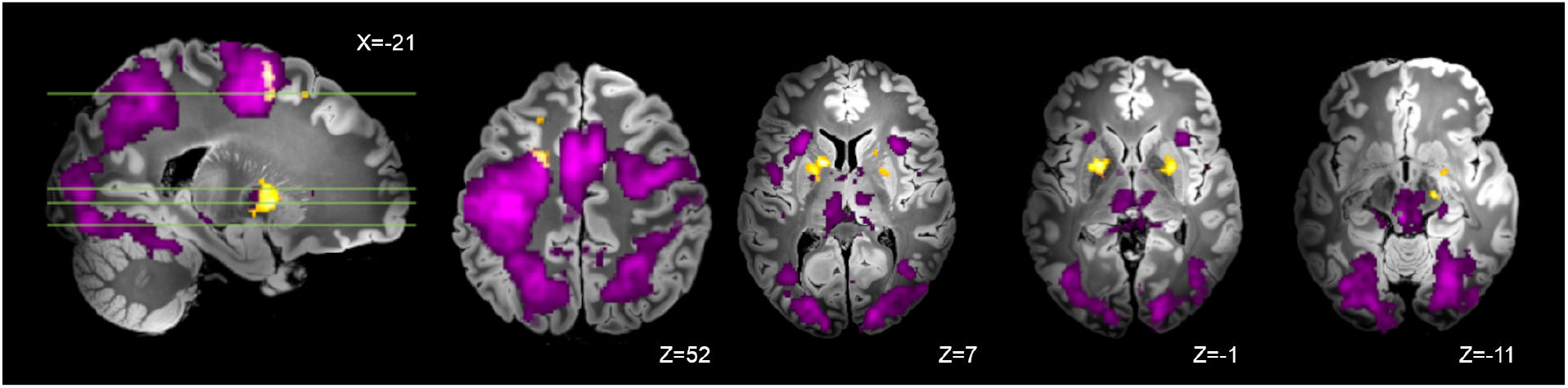
Partial overlap between the differential response to the conditioned inhibitor at training versus test (AX- > DX-, yellow) and the response to the conditioned threat alone (A+, purple) with activation of the left premotor cortex.

